# miRbiom: Machine-Learning on Bayesian Causal Nets of RBP-miRNA interactions successfully predicts miRNA profiles

**DOI:** 10.1101/2020.06.18.156851

**Authors:** Upendra Kumar Pradhan, Nitesh Kumar Sharma, Prakash Kumar, Ashwani Kumar, Sagar Gupta, Ravi Shankar

## Abstract

Formation of mature miRNAs and their expression is a highly controlled process. It is very much dependent upon the post-transcriptional regulatory events. Recent findings suggest that several RNA binding proteins beyond Drosha/Dicer are involved in the processing of miRNAs. Deciphering of conditional networks for these RBP-miRNA interactions may help to reason the spatio-temporal nature of miRNAs which can also be used to predict miRNA profiles. In this direction, >25TB of data from different platforms were studied (CLIP-seq/RNA-seq/miRNA-seq) to develop Bayesian causal networks capable of reasoning miRNA biogenesis. The networks ably explained the miRNA formation when tested across a large number of conditions and experimentally validated data. The networks were modeled into an XGBoost machine learning system where expression information of the network components was found capable to quantitatively explain the miRNAs formation levels and their profiles. The models were developed for 1,204 human miRNAs whose accurate expression level could be detected directly from the RNA-seq data alone without any need of doing separate miRNA profiling experiments like miRNA-seq or arrays. A first of its kind, miRbiom performed consistently well with high average accuracy (91%) when tested across a large number of experimentally established data from several conditions. It has been implemented as an interactive open access web-server where besides finding the profiles of miRNAs, their downstream functional analysis can also be done. miRbiom will help to get an accurate prediction of human miRNAs profiles in the absence of profiling experiments and will be an asset for regulatory research areas. The study also shows the importance of having RBP interaction information in better understanding the miRNAs and their functional projectiles where it also lays the foundation of such studies and software in future.

## Introduction

The regulatory small RNAs in eukaryotes, commonly called as miRNAs, are supposed to regulate at least one third of the genes post-transcriptionally [1]. There are more than 2,000 miRNAs reported alone for human in miRBase and total miRNAs >38,000 [2]. miRNAs are highly spatio-temporal and stand as a cornerstone to a wide range of cellular processes. In general, unlike regular other RNAs whose expressions are mainly controlled at transnational level, miRNAs biogenesis is an event where post-transcriptional regulation plays the most important role in its expression [3, 4]. The highly spatio-temporal formation and expression of miRNAs remains an enigma which traditional canonical model sticking all reasoning on just handful of uniformly expressed RNA Binding Proteins (RBPs) like Drosha/DGCR8/Dicer appear insufficient to answer this [3,5–6].

For a longtime this has been argued that the RBPs like DGCR8, Drosha, and Dicer are indispensable components in miRNA biogenesis. Though these RBPs express themselves in almost all tissues, the miRNAs are mostly highly spatio-temporal and specific in their expression. There are number of regulatory sRNAs whose biogenesis involves many other RBPs besides these regular RBPs. In 2008, a long pending puzzle of regulation of *let*-7 miRNAs was resolved where it was found that RBP LIN28 interaction with *let*-7 miRNA blocked its maturation despite of it being transcribed [7]. By the end of year 2012, a very interesting work with pri-miRNA transcription identified that FUS/FET proteins co-transcriptionally bound some pri-miRNAs, which in turn faciliated the binding of Drosha/DGCR8 complex while influencing the process of miRNA biogenesis [8]. Some recent studies have provided evidences that Argonautes also play a role in miRNA formation in Dicer independent manner [9]. Besides this, a good number of miRNAs have been found originating directly as a product of splicing, called as mirtrons [10]. Some timely reviews have also discussed the involvement of other RBPs in miRNA biogenesis and highlighted the need to focus on this area of research [11].

In 2011, while characterizing miRNAs for their regulatory properties, our group had reported that miRNAs precursors host a number of different RBP binding spots which may control miRNA biogenesis [12]. At that time, due to the scarcity of high-throughput data for cross-linking, the study was limited. Attributable to recent advancements caused by Next Gen Sequencing (NGS) methods, transcriptome wide RBP interactome can now be discovered. These methods include sRNA-seq, RNA-seq, and cross-linking based sequencing techniques like CLIP-seq. In the last decade, volume of CLIP-seq data has increased vastly, which clearly underline the growing acknowledgment for RBPs led regulation in cell system. If the data from sRNA-seq, RNA-seq, and CLIP-seq are considered in a relevant and condition specific manner, an enormous level of information regarding miRNA:RBP regulatory system could be revealed. This would also help to resolve the dynamic and spatio-temporal nature of miRNA biogenesis and build models which could accurately predict the mature miRNA turnover just by using the network information of post-transcriptional regulators.

More than 1,500 RBPs have been identified in human system and this number has been been showing only incremental trends [13]. New RBPs are being identified and new high-throughput cross-linking based studies are revealing the RBP interaction repertoire with every new study. These high-throughput studies have provided a gateway to look into the miRNA and RBP interactions also., Several new RBPs have been found interacting with miRNA precursors. In this direction very recently, two important studies were carried out which deserve special mention. The first one was carried out through proteomics based pull down approach to identify interactions of various RBPs with 72 different human miRNA precursors [14]. This study is one of the first high-throughput studies which provided the direct evidence of several other RBPs beyond Drosha/Dicer interacting with miRNA precursors directly in functional manner. Very next year, a study was reported which used publicly available eCLIP-seq data from two cell lines (HepG2 and K562) from ENCODE and looked for their reported interaction coordinates for overlaps with miRNA regions, followed by experimental validation of 10 RBP-miRNA interactions which directly implicated them involved in miRNA biogenesis [15]. Simultaneously, the authors also observed that miRNA-RBP interactions appeared highly specific to cell types even when studied for just two cell lines conditions. Many RBP-miRNA interactions which were found in one cell line were not present in another cell line despite of expression of RBPs, clearly suggesting that RBP-miRNA interactions varied across the cell states or experimental conditions and other deciding factor may be involved in determining RBP-miRNA interactions and its impact on miRNA biogenesis level. Similar views were echoed by the earlier work of *Treiber et al* which also found that miRNA biogenesis pattern and involved partners varied across the considered cell lines.

What these studies have not answered so far is to explain the spatio-temporal nature of miRNA biogenesis and its causality frame. They have reported some new RBPs found regulating miRNA formation while observing that the interaction varied a lot in their considered cell lines, suggesting the possibilities of multiple paths to regulate miRNA biogenesis. Though, they confirmed the functional associations of some of these identified interactions for a limited number of RBP-miRNA pairs across a limited number of cell lines, it is understandable that validating every CLIP-seq found interaction for larger space through experiments is difficult. These studies did not attempt to infuse other high-throughput data from other platforms like RNA-seq and miRNA-seq to build the reasoning circuits to explain miRNA biogenesis through some models. But these studies strongly pointed out the need to provide the explanations for the observed spatio-temporal nature of miRNA biogenesis and strongly advocated the need to look with bigger picture of interactions which our present study has taken up. Such reasoning threads would be highly important to explain as well as predict miRNA formation for wide range of conditions. In such scenario, the computational modeling of these interactions becomes an urgent need. Such computational studies can explore and discover much larger horizon of interactions having functional validity as well as discover the causal circuits associated with these interactions which could explain the spatio-temporal nature of miRNAs for much larger space and extent of universality.

In the present work, we have carried out the study across a large number of experimental conditions (47) while having >25 TBs of data covering high throughput experimental data from various CLIP-seq platforms, sRNA-seq, and RNA-seq data. Considering these data together may help to build the causal networks and logical units which could reason such spatio-temproal interactions and their impact on miRNAs biogenesis level. These conditional causal networks explaining the miRNA biogenesis were identified using Bayesian network analysis (BNA) which were finally molded into a machine learning system, XGBoosting, to accurately predict miRNAs profile for any given condition. It was achieved for 1,204 miRNAs. This system is capable to give an accurate prediction of miRNAs profiles in the absence of miRNA profiling experiments like miRNA-seq or array experiments while building upon the identified miRNA-RBP specific Bayesian network (BN) components dynamics and information in the back-end. A first of its type, this tool has been tested across a large number of experimentally validated data from a wide range of experimental conditions and cell types where it displayed consistently high accuracy and reliability.

## Methods

### Source of NGS Data and Data processing

Data for sRNA-seq and RNA-seq based high throughput studies were collected from Gene Expression Omnibus (GEO) and Sequence Read Archive (SRA) for 21 experiments (data volume ~15.6TB) which included 47 different experimental conditions. Complete list of various experimental conditions (for both RNA-seq and sRNA-seq) and source of data along with detailed experimental description are available in Table 1. The miRNA sequences of human (1,881 pre-miRNAs, 2,588 mature miRNAs) were collected from miRBase database version 21 [2]. 1,230 CLIP-seq experimental samples were collected for 155 RBPs from ENCODE and GEO database (~10.8 Tb). These data were derived from different types of CLIP-seq experimental techniques (PAR-CLIP, HITS-CLIP, eCLIP, iCLIP, CLEAR-CLIP and irCLIP). The detailed description of CLIP-seq datasets is given in S1 Table.

**Table 1.** This table describes the different RNA-seq and sRNA-seq experimental data collected in the current study.

| RNA-seq |  | sRNA-seq |  | Experiment | Conditions |
| --- | --- | --- | --- | --- | --- |
| Accession no. | Study id | Accession no. | Study id |  |  |
| GSE56862 | SRP041228 | GSE56862 | SRP041228 | PolyA independent deep sequencing of the chromatin-isolated RNA fraction. | Cervical normal |
| GSE63511 | SRP050087 | GSE63511 | SRP050087 | Thyroid tissue | Thyroid tumor , Thyroid normal |
| GSE68631 | SRP058087 | GSE68631 | SRP058087 | HEK293 cells in 6-well plate were transiently transfected with 400 ng plasmids of control, shRNA, shRNA-LC, siRNA-RZ for 48 hr. | HEK293 cell |
| GSE69446<br>& | SRP058953<br>& | GSE69446<br>& | SRP058953<br>& | Healthy controls and Crohn's patients | Monocytes_chrone,<br>Monocytes_normal |
| GSE66209 | SRP055438 | GSE66209 | SRP055439 |  |  |
| GSE69787 | SRP059380 | GSE69787 | SRP059380 | Ossification of the posterior longitudinal ligament (OPLL) tissue and normal posterior longitudinal ligament (PLL) tissue | OPLL and PLL. |
| GSE86491 | SRP090091 | GSE86491 | SRP090091 | Endometrial tissue from two time points of the menstrual cycle | Endometrial_TP1, Endometrial_TP2 |
| GSE37764 | SRP012656 | GSE37764 | SRP012656 | Primary non-small cell lung adenocarcinoma tumors and normal tissues. | Lungs Normal , Lungs Tumor |
| GSE67491 | SRP056784 | GSE67491 | SRP056785 | Whole blood samples personalized medicine study | Caucasian male blood,,Caucasian female blood, African_america_female_blood |
| GSE70666 | SRP060566 | GSE70666 | SRP060565 | Oral squamous cell carcinoma | Oral_cancer |
| GSE85145 | SRP080859 | GSE85145 | SRP080860 | Primary Human Astrocytes Infected with Borrelia burgdorferi | Astrocytes_infection |
| ERS327013 | ERP003613 | SRP023531 | GSE47720 | Human pancreatic islets and enriched beta-cells | Pancreatic beta-cells |
| GSE71336 | SRP061565 | GSE71336 | SRP061566 | LNCaP cells expressing the wild-type androgen receptor (AR-WT) or the ligand-independent AR-V7 splice variant | Lncap cell |
| GSE65515 | SRP053046 | GSE65515 | SRP053046 | To reveal dynamic changes in networks of gene expression and epigenetic regulation during healthy human T cell aging. | Newborn, Middle-aged, long-lived |
| GSE92876 | SRP095604 | GSE92874 | SRP095605 | Examination, identification and comparison of mRNA expression profiles in control and schizophrenia npc. | Disease and control |
| GSE79032<br>&<br>GSE62830 | SRP071331<br>& SRP<br>049391 | GSE79032<br>&<br>GSE62830 | SRP071334<br>&<br>SRP049389 | RNA Sequencing Reveals that Kaposi Sarcoma-Associated Herpesvirus Infection Mimics Hypoxia Gene Expression Signature & Differential Expression Profiles of miRNA- mRNA Target Pairs in KSHV-Infected Cells | Infected, KSHV-infected |
| GSE73502 | SRP064515 | GSE73502 | SRP064235 | Widespread shortening of 3' untranslated regions and increased exon inclusion characterize the human macrophage response to infection | Salmonella, Non-infected, Listeria ( each condition in 2hrs and 24 hrs) |
| GSE78169 | SRP070657 | GSE78169 | SRP070659 | An integrative transcriptomics approach identifies miR-503 as a candidate master regulator of the estrogen response in MCF-7 breast cancer cells | Breast cancer at 10 different time point. |
| GSE46224 | SRP021193 | GSE46224 | SRP021193 | Dynamic regulation of myocardial noncoding RNAs in failing human heart and remodeling with mechanical circulatory support | Heart_ischemic, Heart_nonfailing, Heart_nonischemic |
| ERS327018 | ERP003613 | SRX262197 | SRP018255 | Placenta normal tissue | Placenta |
| ERS326957 | ERP003613 | SRX271415 | SRP021475 | Testis normal tissue | Testis |

It provides the details of accession ID, study ID and different experimental conditions considered in this study.

The RNA-seq data was filtered using filteR [16] ensuring that the quality value of at least 70% length of read should have scored QV30. filteR has inbuilt adapter removal module and cleans the read data for errors. sRNA-seq data was processed using trimmomatic [17]. Same quality filtering measures were taken while considering the minimum cut-off read length of 18 bases. For mapping RNA-seq reads across the human genome build 38 assembly (hg38) Seqmap was used which is based on Bowtie platform [18]. The alignment results were saved in SAM format for expression quantification. rSeq was used for quantification of gene expression from SAM files [19]. Normalized expression values like RPKM/FPKM were considered for RNA-seq and TPM was considered for sRNA-seq. For expression analysis of pre-miRNAs, same RNA-seq pipeline was used where RNA-seq reads were mapped over known pre-miRNA sequences collected from miRBase. The alignment results were stored in a SAM file. rSeq was used for quantification of pre-miRNA expression from RNA-seq data. miRDeep2 was used for expression analysis of mature miRNAs from sRNA-seq data with mismatch value of “2” [20]. For mapping sRNA-seq reads across the known mature miRNAs, miRDeep2’s mapper.pl script was used. miRDeep2’s quantifier.pl script was executed for the quantification of mature miRNAs expression from the saved SAM files.

Before performing the further analysis, the RNA-seq and sRNA-seq data were corrected by removing the batch effect. Batch effect correction is the procedure of removing variability from data that is not due to variable of interest but arises due to technical differences between samples i.e. the type of sequencing machine, different experimental conditions or even the technician that ran the sample. The batch effects were corrected for each sample using the limma R-package [21] considering a two-way ANOVA approach. The Limma package fits a linear model containing a blocking term for the batch structure to the expression values for each gene. Subsequently, the coefficient for each blocking term was set to zero and the expression values were computed from the remaining terms and residuals, yielding a new expression matrix without the batch effects.

Out of 155 RBPs, CLIP-seq data of 46 RBPs were collected from ENCODE database. CLIP-seq data for 109 RBPs were collected from GEO database. All these raw reads were processed following a standard protocol. In this study only those RBPs were considered which had at least two samples reported. Out of 155 RBPs, 12 RBPs data were from studies done on only one sample. These RBPs were discarded from the current study. For considering the most probable binding sites of RBPs on different miRNAs a criteria was set that at least five reads should map to the considered region of miRNA and at least two different samples must support it (Approach#1). The binding sites qualifying this step were further screened through two peak calling tools, Piranha (Approach#2) and PEAKachu (Approach#3) (https://github.com/tbischler/PEAKachu) [22]. Both tools were run with default parameter settings. Piranha was run using ‘piranha -s -a 0.95 <.bed>’. PEAKachu requires control file in native mode, therefore, it was run in adaptive mode using ‘peakachu adaptive -t <.bam>’ where it uses certain amount of dataset from given file as control to find the peak, concurrence of Approach#1 with at least any one of these two tools (Approach#2/Approach#3) was made mandatory to consider the binding sites valid for further downstream analysis.

pre-miRNAs information was collected from the hairpin loops provided at miRBase database. Also these sequences were extended up to 1kb flanking regions on both the sides. Such regions were considered as putative pri-miRNAs in our study which essentially covered highly probable pri-miRNA regions [23]. For processing of the CLIP-seq data FastX (http://hannonlab.cshl.edu/fastx_toolkit/) was used. The adapters reported in literature for respective CLIP-seq data were used for filtering. To remove low quality reads, only those reads were kept which had at least 75% of bases with a quality score of 25 or more (-Q33 -q 25 -p 75). Unique reads were selected. The unique CLIP-seq reads were mapped across pri-miRNA and pre-miRNA regions using Bowtie while considering a maximum of two mismatches to get the possible binding sites (-f - S -n 2 -a).

### Expression and network analysis

CLIP-seq data and RNA-seq data having same experimental conditions were collected from GEO and ENCODE databases. The CLIP-seq data collected for 138 RBPs had 82 experimental conditions. The data collected for RNA-seq and sRNA-seq covered 47 experimental conditions. Total 32 experimental conditions were common among all the three types of high-throughput data. Possible interacting partners were collected for each RBP from STRING database [24]. Maximum up to eight steps of interactions were considered for the primary network construction. S1 Table provides complete details on the binding sites data collections. S2 and S3 Table provides complete details on the TCGA data collections for model building and validation and Table 1 provides complete details on the expression data of sRNA-seq as well as RNA-seq used in the present study. Data for cancer specific cluster analysis using miRNAs and associated RBPs were taken from GEO with accession IDs: SRP012656 and SRP050087.

### Reconstruction of miRNA:RBP association using Bayesian network analysis (BNA)

BNA was performed for the considered experimental conditions separately. Its input consisted of the expression data of pre-miRNAs, mature miRNAs, RBPs, and associated genes obtained from their respective PPI data. Those RBPs having binding sites across those expressed miRNAs (including pri/precursors) for the given experimental condition were considered in the BNA. Any particular RBP in the pre-miRNA-RBP interaction modeling was included if it had binding sites in the primary and pre-miRNAs regions. Similarly, any particular RBP was included in the mature miRNA:RBP interaction modeling if it had binding site in the pre-miRNA sequence for the corresponding mature miRNA. This analysis was followed for each experimental condition separately and the miRNA:RBP associations were obtained for the expressed miRNAs for both the major steps in its biogenesis (from pri-miRNA to pre-miRNA, and pre-miRNA to mature miRNA processing steps).

Since RBPs and associated PPI components are the causal factors of miRNA processing, a more general approach was adopted via structural equation modeling (SEM) [25, 26]. It has two parts: one is response (dependent variable) and another is independent variables. The basic idea in SEM is to estimate the potential RBPs involved in each step of miRNA biogenesis. As our data (expression data of RBP, associated genes from PPI, pre-miRNA, and mature miRNA) are continuous in nature, it was assumed as a multivariate Gaussian distribution for all nodes throughout the study. The basic model used in the current study is a p-dimensional random vectors X= (X_1_,X_2_,…, X_p_) with joint distribution P(X_1_,X_2_,…, X_p_). Here, the variable X_1_,X_2_,…, X_p_ are the nodes in the network which represent RBP, associated genes from PPI, pre-miRNA, and mature miRNA. BNs are directed graphical model and their edges encode conditional independence constraint implied by the joint distribution of **X**, which is:

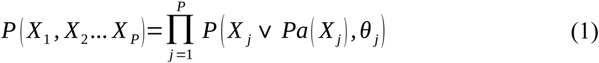

Here, Pa (*X_j_*)=[*X_i_*: *X_i_→ X_j_∈E*] is the parent set of *X_j_* and *θ_j_* encode parameters that define the conditional probability distribution (CPD) for *X_j_*. The different steps followed for the construction of miRNA biogenesis model are as following:

1) Estimation of Directed Acyclic Graphs (DAGs) between miRNA and RBPs, RBPs with respective PPI partners based on expression data of miRNAs, RBPs, and their associated proteins. The binding sites of RBPs on miRNA were considered as the prior information.

2) Identification of significant DAGs between miRNAs and RBPs, RBPs with their respective PPI partners were taken out from the total DAGs considering a suitable convergence criteria (Error tolerance < 10^−4^) in block coordinate descent algorithm [27].

3) It was assumed throughout the study that the data were generated from a multivariate Gaussian distribution, where error covariance matrix is a positive definite. Thus, the significant DAGs obtained in the previous step were directly modeled through a generalized linear model.

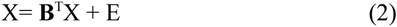

This is called a SEM for total observations **X**. **B** is the weighted adjacency matrix of a directed graph and E ~ N (0, w_j_^2^).

4) The significant DAGs were modeled using SEM. To avoid the high-dimensional data structure (i.e. n<<p), sparse regularized penalty was used [28]. Following model was considered for the data:

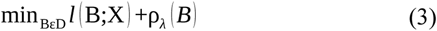

where, *D ⊂ R^p × p^* is the set of weighted adjacency matrix that represents a acyclic directed graphs for all nodes (significant and non-significant) from all the three terms: the loss function l, the constraint D, and the sparse regularizer *ρ_λ_*. Whereas, B is a weighted adjacency matrix for significant edges *BεD*.

5) In this study, to fit the model an array of penalties (λ ranging from 1-10) was decided and an optimum penalty was selected which fitted to the data using the minimax concave penalty (MCP) algorithm [29]. Method of least squares regression was used to regress between node and its parent as the data is continuous to estimate the parameters.

6) Further, to improve the estimation of parameter, weighted adjacent matrix was represented by B as B=est ( *β*) which was used to estimate the conditional variance by the given formula:

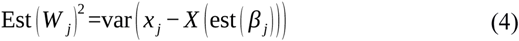

*Ω* =diag (est (*W*_1_), …,est (*W_p_*)) was applied as a variance matrix and combining [est (*B*), est (*Ω*)] to calculate the variance covariance matrix Σ and accuracy for each parameter. A suitable convergence criteria (Error tolerance < 10^−4^), precision value >=85% and an alpha threshold of 0.05 were considered for the selection of the parameters. These parameters decided how larger the effect size (positive/negative) was between miRNA and RBP.

The algorithm followed in BNA was performed using “Sparsebn”, “ccdr Algorithm” and “SparsebnUtils” R-packages. The associated basic steps followed in the current approach are described in Fig 1.

**Fig 1.**
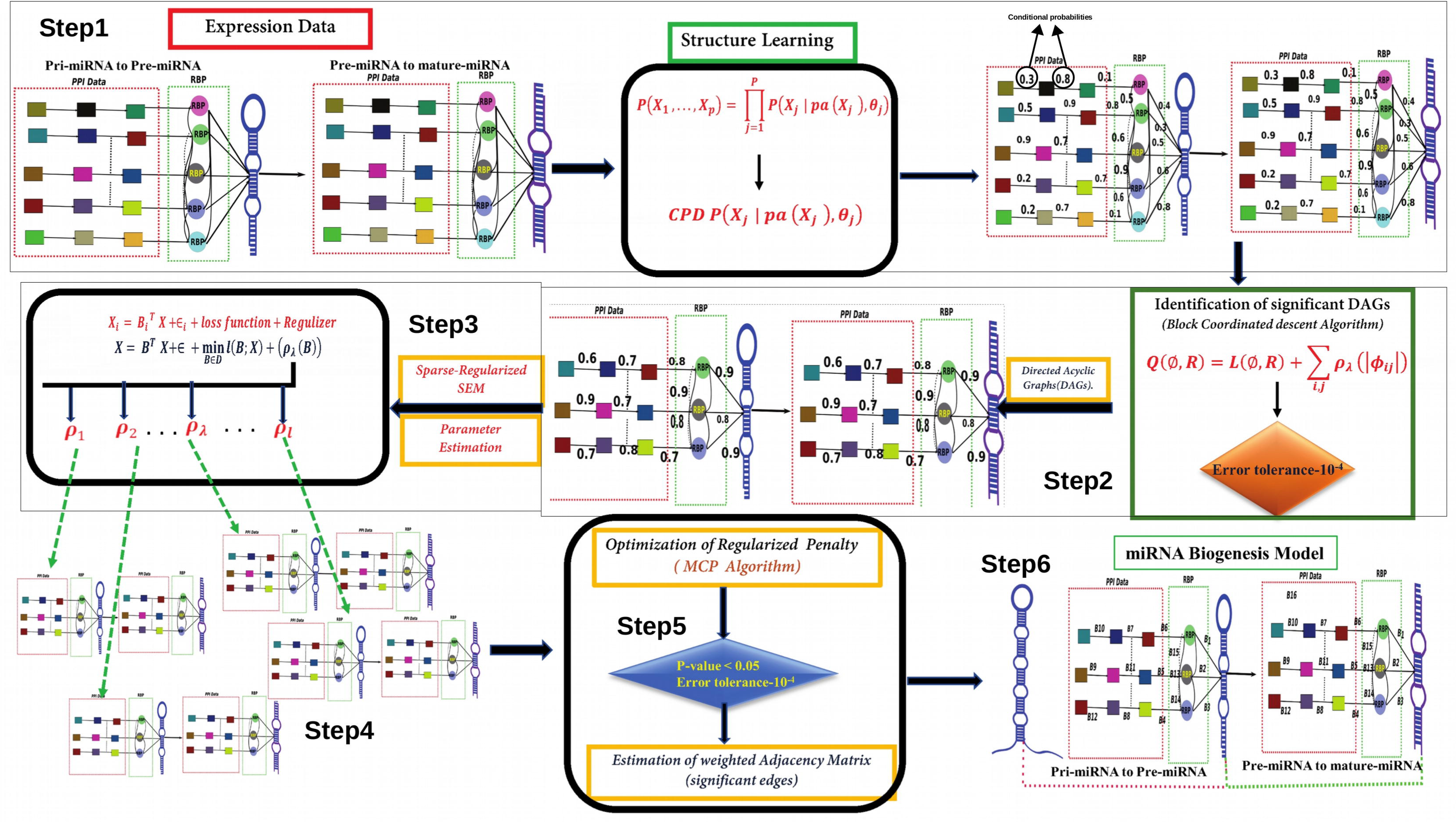
The basic steps taken in Bayesian network reconstruction. Different steps were followed as: 1) Estimation of DAGs between RBP and miRNA, RBP and its PPI partners based on the expression data and prior information. 2) Identification of significant DAGs from total DAGs considering a suitable convergence criteria. 3) The significant DAGs obtained in previous step were directly modeled via a generalized linear model, assuming a multivariate Gaussian distribution. 4) Parameter estimation between RBP and miRNA, RBP and its PPI partners based on the significant DAGs using a sparse regularized penalty. Different set of rho lambda values were taken into consideration and optimum lambda value were selected. 5) Selection of optimum sparse regularized penalty from an array of penalties that fit best to data. 6) Final estimation of parameters using the optimum penalty and selection of parameters considering certain suitable criteria.

### Functional assessment of miRNA:RBP associations

From the result of BNA, miRNA:RBP combinations were collected. RBPs appearing responsible (positively/negatively associated based on p-value (alpha-threshold) <0.05) in the processing of pri-miRNA, and mature miRNA along with their respective back-chains in each experimental condition were collected. The association of RBPs with miRNAs (pre-miRNA and mature miRNA) was verified based on the expression correlation coefficient cut-off of >=|0.6| as this is considered as a good cut-off value for such studies [30] RBP expected to be involved in processing a pre-miRNA from its pri-miRNA exhibits a positive correlation with the corresponding pre-miRNA. Similarly, the RBPs apparently involved in mature miRNA formation were supposed to display positive correlation with mature miRNA and negative correlation with its pre-miRNA.

The miRNA:RBP associations obtained in this study were validated across eight different independent normal tissues (bladder, testis, brain, breast, lungs, pancreas, placenta and saliva), offering totally different source of testing data. The mature miRNAs expression data were collected from miRmine database [31] and RNA-seq expression data were collected from GTEx, ARCHS4 [32], and Array Express [33]. Correlation analysis was performed between mature miRNA expression and RNA-seq expression data. Batch effect normalization was followed as described above.

To evaluate the functional association of miRNAs and their associated RBP, a functional enrichment analysis was performed. The mature miRNAs were clustered based on their expression data (combining the eight tissues) and those RBPs associated with the cluster of miRNAs were studied for RBPs common among the group of miRNAs in any given cluster. Those mature miRNAs were considered for clustering where at least 50% samples exhibited their expression. A functional enrichment analysis (for pathway, molecular function and biological process) was performed using Enrichr [34] for miRNA targets and the associated RBPs for each cluster of miRNAs having an FDR <= 0.05. Experimentally validated miRNA targets were collected from miRTarBase [35]. The common pathways, molecular functions, and biological processes were checked for each miRNA cluster and their associated RBPs.

### RNA-seq based potential miRNome profile detection using XGBoost regression

In-house developed python programs were used to carry out the machine learning exercises. Python scikit-learn and keras libraries were used. The objective was to build the predictive models of miRNA expression based on the RBP’s network components’ expression data. For the prediction of miRNA expression level the interaction network of RBPs and their associated genes obtained from the BNA were used. For every given miRNA, its associated Bayesian network genes were considered as the input feature variable which carried their values of expression level for any given condition while the associated miRNA’s expression in that condition worked as the target value. To build the predictive model, XGBoost regression was applied [36]. XGBoost is an extreme gradient boosting ensemble technique where prediction is done by an ensemble of simple estimators giving higher weights on difficult learning instances to minimize the loss function using gradient. For a given dataset with *n* samples and *m* features D= {(*Χ_i_*, y*_i_*)}(|*D*|=n,X_*i*_ɛR^*m*^, y*_i_*ɛR), an ensemble tree model uses *K* additive functions to predict the output:

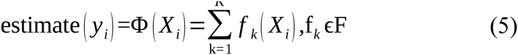

where F=f*_x_*=w_*q*(*x*)_(*R_m_*→ T,WɛR^*T*^) is the space of regression trees. Here ‘***q***’ defines the structure of each tree that maps its corresponding leaf index. ‘T’ is the number of leaves in a tree. Each *f_k_* corresponds to an independent tree structure q and leaf weight w. Unlike decision trees, each regression tree contains a continuous score on each of leaf, we used *w_i_* to represent score on *i*-th leaf.

To learn the set of functions used in the model, we minimized the following regularized objective:

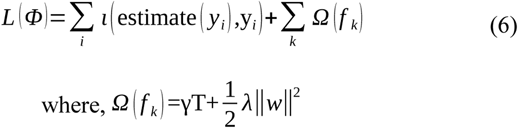

Here ‘***t*’** is a differentiable convex loss function that measure the difference between the prediction estimate (*y_i_*) and target *y_i_*. The second term, **Ω**, penalizes the complexity of the regression tree function. The additional regularizer term helps to smoothen the final learned weights to avoid over-fitting. The regularized objective finally selects a model applying the predictive functions.

**Datasets:** The RNA-seq and sRNA-seq expression data were collected from TCGA database for seven different tissues (both normal and cancer). These were independent from the above described 47 experimental conditions used in the construction of Bayesian Nets for miRNA biogenesis. For model building only those miRNAs were considered which had sufficient number of instances to train and build i.e. only those miRNAs were considered for which at least 600 sample evidences were present for expression. This way, models for 1,204 miRNAs could be formed. A total 17,737 of RNA-seq samples and 11,717 miRNA-seq samples, covering 30 experimental conditions and 15 tissue types, were used to train and build the models. As per the default practice in machine learning training, 70% of the dataset was used to train the XGBoost system while testing was done over remaining 30% data for each miRNA. 10 fold Cross Validation analysis was also done during the training step for every miRNA to check the consistency of the model.

For further additional validation, other eight different tissues’ conditions (liver, postrate, pancreas, ovary, lungs LAUD, Lungs LUSC, adrenal gland ACC and adrenal gland PCPG) data were used separately from TCGA, covering 431 samples. Full details about these training, testing, and re-validation data are given in S2 and S3 Table.

The predictive models were validated considering the following statistical measures:

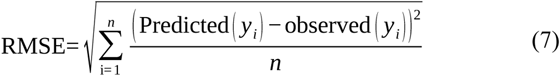

To evaluate the predictive accuracy of each model, Relative Mean Absolute Percentage Error (RMAPE) was used. The RMAPE is the prime metric used widely to validate forecast accuracy, providing an indication of the average size of the prediction error expressed as a percentage of the relevant observed value. The equations for RMAPE and accuracy are given below as:

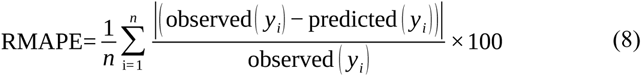

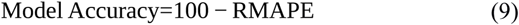

## Results and Discussion

Canonical model of miRNAs biogenesis is usually classified into a two steps process where in the first step the transcribed pri-miRNA is processed into pre-miRNA and in the second step the mature miRNA is formed from the pre-miRNA. In this study 1kb flanking regions from both sides of published pre-miRNA sequences were considered as putative pri-miRNA as there is barely any resource available for pri-miRNA identification. Considering 1 kb flanking region is based on well practiced protocol while considering the fact the pri-miRNAs size starts with minimum ~1 KB [23]. This also ensured that RBP interacting closer to precursor releasing sites would be covered reasonably. To find out the possible interactions with RBPs, these miRNAs were first scanned for RBP bindings using publicly available CLIP-seq data which were mapped across the miRNA sequences (covering primary, precursor, and mature miRNA regions) and downstream studies were carried out at different levels.

### CLIP-seq data analysis provides the most potential candidate RBPs to start with to build the RBP-miRNA causal nets

A total of 1,230 CLIP-seq samples for 155 RBPs were obtained from ENCODE and GEO, covering 10.8 TB reads. These reads were checked and filtered for quality and were mapped across the considered sequences of each known human miRNA using the protocol described in the methods section. Those interactions were considered as the valid starting points for downstream analysis where at least five reads mapped in at least two different samples (Approach#1). Further to this, the interactions were evaluated for peak calling using two different tools, Piranha (Approach#2) and PEAKachu (Approach#3). A total of 194,795 and 494,753 peaks were reported by Piranha and PEAKachu, respectively. Both these tools agreed with each other for just 24.76% of their result peaks, suggesting substantially different results reported by these two peak calling tools. Likewise, it was found that they discarded even some potentially true positive cases where the interactions were found repeatedly happening across several different experimental conditions, when considered individually. For example, PRPF8 is reported for processing of precursor to mature miRNA miR-25 and miR-93 [37]. Approach#1 detected these interactions in all the samples of PRPF8 but Piranha failed to capture this. However, their individual agreements with Approach#1 were much higher than among themselves, clearly suggestive of effective screening when these approaches as used in combination. The identified RBP:miRNA interactions screened through Approach#1 overlapped with ~80% of the identified interaction partners by the two peak calling tools whose outcome made the final starting set of most confident RBP-miRNA interactions for the downstream analyses. Thus, a total of 138 RBPs qualified the criteria for interactions with the putative pri-miRNA exclusive regions and 126 RBPs qualified for interactions with pre-miRNA regions for the further downstream analysis. The binding sites were clustered based on RBPs, pre-miRNAs, and pri-miRNAs. Illustration of binding sites distribution for each RBP, pri-miRNAs, and pre-miRNAs is given in Fig 2 (A,B,C) whose details are provided in S4 and S5 Tables. It was interesting to observe that number of binding sites for RBPs across the miRNAs differed a lot among themselves. Total experiments/conditions taken to report a data also influenced this count. Some RBPs have been studied multiple times which make their count higher. However, when these counts were normalized by the total experimental conditions, clearer picture emerged. Fig 2 (C) illustrates the case for average binding sites for RBPs when normalized by different experimental conditions. RBPs like AGO4, HNRNPH1 and FMR1 much higher number of target pre-miRNAs than other RBPs, though the count is much lesser than the not normalized values reported in the bubble diagram. While SRRM4, MBNK2, and PRKRA like RBPs displayed the lowest average binding sites across the pre-miRNAs. On the average there were ~63 pre-miRNA targets per RBP. As Fig 2C suggests, a lot of variations was observed in this value across the RBPs. Also, it was evident that the binding of RBPs varied a lot across the different experimental conditions. Some RBPs showed positional preferences (Fig 2D). RBPs like Argonautes, DGCR8, and PUM2 exhibited preferences for pre-miRNA regions. Interestingly, Drosha displayed almost uniform distribution for the binding sites. Distribution of binding sites per miRNA for each RBP is given in Fig 2(E), which was also found highly variable for each RBP.

**Fig 2.**
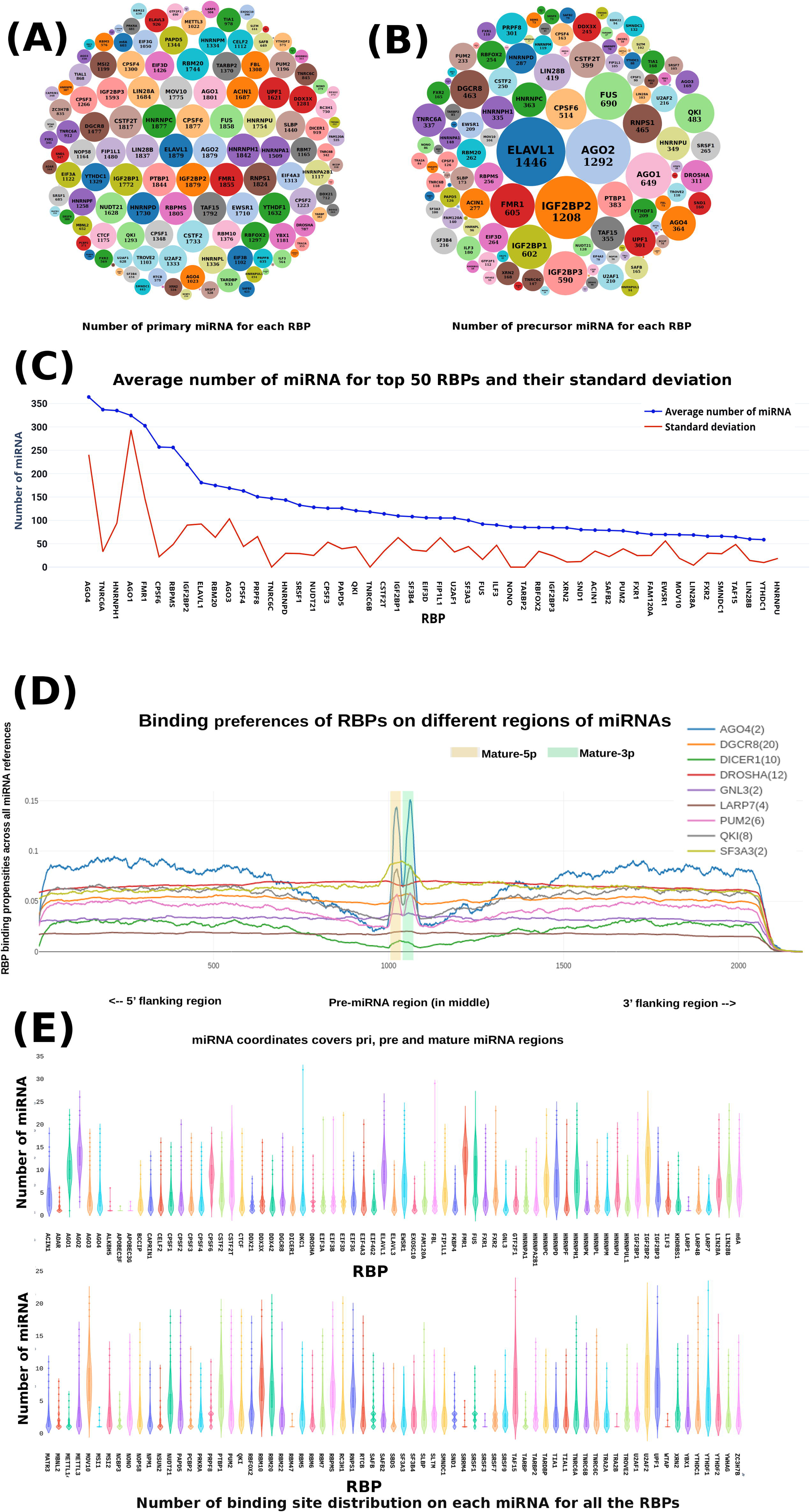
Distribution patterns for RBP binding across miRNAs. A total of 138 RBPs were binding to the primary regions of miRNAs whereas 128 RBPs were binding to the precursor regions of miRNAs. (A) Bubble chart showing number of primary miRNAs bound by each individual RBP. (B) is showing the same for of precursors. (C) Number of unique pre-miRNA targets for various RBPs when normalized by total number of different experimental conditions considered. The total number of unique targets for each RBP increases with the number of experimental conditions considered. Different numbers of studies have been conducted for each RBP, which results into skewed data reporting for total targets if not properly normalized. (D) Binding preference of some RBP where it can be observed that some RBPs prefer to bind in the precursor regions. (E) Violin plot illustrating the distribution of the number of binding sites per miRNA for each RBP.

The CLIP-seq data analysis provided the initial starting point to consider the most potential interacting miRNA-RBP pairs. Implementation of BNA (in the following section) over such interactions further screened such interactions for only those associations which significantly displayed functional association with high confidence while building upon the expression and interaction network information.

### Bayesian networks derive the functional RBP-miRNA associations along with causality routes

The CLIP-seq data provided the first stop to start with where RBPs and miRNAs associations were derived for binding. All such RBP-miRNA associations may not be functional and may not be involved in the miRNA biogenesis. In order to screen for the functional RBP and miRNA associations, a Bayesian network analysis was performed. RBPs interactions and influence on miRNAs may depend upon other interacting factors which could together work in a dynamic network and define causality for miRNA biogenesis. This dynamic causal network was built on the foundation of the primary information of association between the RBPs and miRNAs derived from the CLIP-seq data as mentioned above. To this, the protein-protein interaction data available for the associated RBPs were looked into and fitted for all the available conditions to define the static primary networks. In order to get the directionality/causality, conditional dynamic network was needed. This was achieved by mapping the RNA-seq and sRNA-seq expression data for all the genes and the associated miRNA in the network, for all the available experimental conditions. The methods section has described the steps implemented through the BNA to derive the causal nets.

~97% combinations obtained after the BNA (which included the miRNAs, associated RBPs and their causal network genes in the back-end) displayed very strong correlation (>=|0.8|). A total of 8,047 pre-miRNA:RBP (RBPs supposedly involved in formation of pre-miRNA from pri-miRNA) and 10,100 mature miRNA:RBP (RBPs supposedly involved in processing of mature miRNA from pre-miRNA) unique combinations were obtained considering all experimental conditions together. This covered causal networks for a total of 746 precursors and 1,346 mature miRNAs. These pre-miRNA:RBP and mature miRNA:RBP combinations obtained from BNA were also verified based on expression correlation in each experimental condition. In each experimental condition more than 89% miRNA:RBP combinations (both pre-miRNA:RBP and mature miRNA:RBP) obtained from BNA were found showing similar type of associations (positive/negative) with strong expression correlation for any given experimental condition.

The miRNA:RBP combinations obtained from this network analysis were independently validated across eight different normal tissues viz. bladder, testis, brain, breast, lungs, pancreas, placenta and saliva. The main interest of this validation work was to measure the extent to which the observed behavior of miRNA:RBP associations was reflected across other sets of tissues which were not included earlier in the current study. Here, the study was focused on the mature miRNA:RBP combinations. Correlation analysis was performed between mature miRNA expression and RNA-seq expression. Those combinations obtained for mature miRNA:RBP from the BNA were evaluated across all the eight tissues considering an absolute correlation coefficient cut-off value of >=|0.6| which is considered as a good cut-off value [30, 38].

As transpires from Fig 3, it was found that 96.9% the identified miRNA:RBP combinations from the BNA were present in at least one of these eight tissues. Also, it was noticed that 49% of miRNA:RBP combinations were present in more than four tissues. There were 270 such miRNA:RBP combinations which appeared in all the eight tissues. To check the significant existence of miRNA:RBP combinations obtained in this study, a Fisher’s exact test was performed using a total of 164,662 random miRNA:RBP combinations which worked as the background. The random miRNA:RBP combinations were searched against the considered eight tissues ( bladder, testis, brain, breast, lungs, pancreas, placenta and saliva). Only 144 random miRNA:RBP combinations (0.09%) were found existing across the eight tissues. Fisher’s exact test was performed considering multiple hypothesis testing which was found significant (simulated p-value <0.01; range scale: 0.03 to 10^−7^), implying that the observed miRNA:RBP combinations across the eight different independent normal tissues were not random at all. Also, each combinations obtained in this study were tested separately using a binomial test against the miRNA:RBP combinations in the random dataset. Out of 10,100 miRNA:RBP combinations 9,786 combinations (96.9%) were found significantly existing (p-value <0.01). From the published literature, 104 experimentally validated functional combinations of miRNA:RBP were obtained as a test set (Table 2). A total of 85 out of 104 such experimentally validated combinations were present in the above mentioned combinations detected by the BNA. This highlighted the effectiveness of the BNA in deriving the functional RBP-miRNA associations and reliability of the large number of novel combinations of miRNAs, RBPs, and causal networks detected by the BNA.

**Fig 3.**
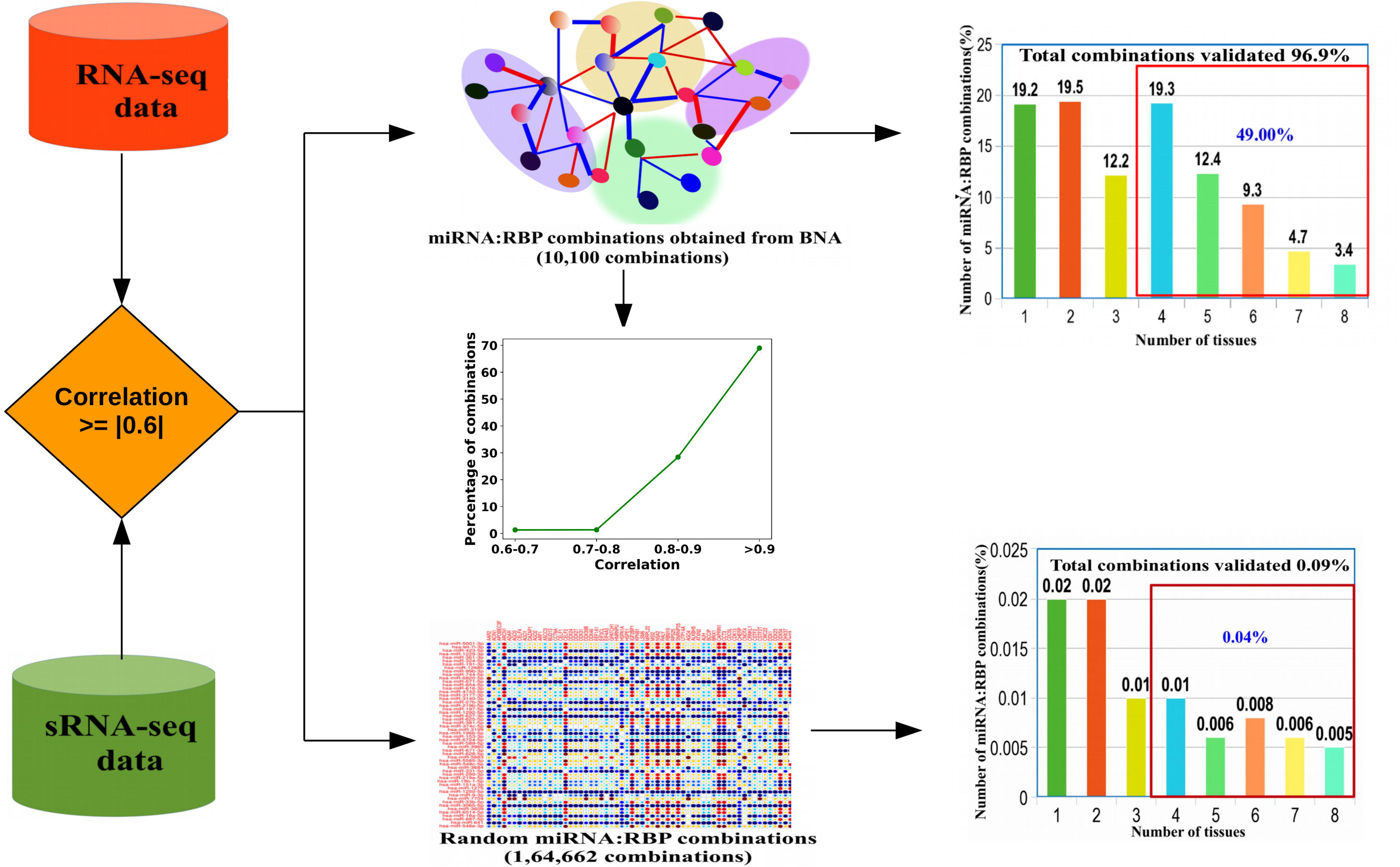
Validation of different miRNA:RBP combinations across totally different eight normal tissues (Bladder, Brain, Breast, Lungs, Pancreas, Placenta and Testis). The bar-chart shows the distribution of unique miRNA:RBP pairs across different tissues. A total of 96.9% miRNA:RBP combinations were validated across all the eight tissue which were never considered before while dealing with data or reconstructing the networks. 49% of miRNA:RBP combinations exist in four or more tissue. Only 0.09% of random miRNA:RBP combinations were found in them whose just 0.04% combinations existed in four or more tissue. Fisher exact test between the distributions for combinations obtained from BNA and randomized combinations was found significant. Also, all one-to-one binomial tests were found significant (p<<0.05) between the BNA obtained and randomly obtained combinations.

**Table 2.** Experimentally validated miRNA: RBP associations collected from various literatures.

| **RBP** | **miRNAs (Pri-miRNA to Pre-miRNA)** | **Association** | **References** |
| --- | --- | --- | --- |
| HNRNPA1 | hsa-mir-18a | positive | [39] |
| FUS | hsa-let-7a, hsa-let-7f, hsa-mir-149, hsa-mir-186, hsa-mir-5001, hsa-mir-636, hsa-mir-652, hsa-let-7b, hsa-mir-103, hsa-mir-106a, hsa-mir-124, hsa-mir-132, hsa-mir-135b, hsa-mir-143, hsa-mir-146a, hsa-mir-149, hsa-mir-16, hsa-mir-182, hsa-mir-18a, hsa-mir-191, hsa-mir-192, hsa-mir-194, hsa-mir-197, hsa-mir-199a, hsa-mir-19a, hsa-mir-19b, hsa-mir-218, hsa-mir-21, hsa-mir-22, hsa-mir-23b | Positive | [8] |
| HNRNPA2B1 | hsa-mir-103a, hsa-mir-3651, hsa-mir-6516 | Positive | [40] |
| METTL3 | hsa-mir-4284 | Positive | [41] |
| SRSF1 | hsa-mir-7, hsa-mir-221 | Positive | [41] |
| DDX3X | hsa-mir-20a, hsa-mir-1, hsa-mir-141, hsa-mir-145, hsa-mir-19b, hsa-mir-34a | Positive | [42] |
| **RBP** | **miRNAs (Pre-miRNA to mature miRNA)** | **Association** | **References** |
| ILF3 | hsa-miR-144 | Positive | [15] |
| RBFOX2 | hsa-miR-144, hsa-miR-18a, hsa-miR-126 | Positive | [15] |
| SF3B4 | hsa-miR-4745, hsa-miR-6753 | Positive | [15] |
| AUF1/HNRNPD | hsa-miR-122 | Negative | [43] |
| CPSF1 | hsa-miR-17, hsa-miR-19a, hsa-miR-20a, hsa-miR-19b | Positive | [44] |
| DDX3X | hsa-miR-122 | Positive | [45] |
| DKC1 | hsa-miR-664 | Positive | [46] |
| AGO1 | hsa-miR-30a, hsa-miR-182, hsa-miR-21, hsa-miR-183, hsa-let-7f, hsa-let-7a, hsa-miR-30a, hsa-miR-378a, hsa-miR-140, hsa-miR-92a | Positive | [47] |
| AGO2 | hsa-miR-451, hsa-miR-30a, hsa-miR-21, hsa-miR-92a, hsa-miR-99b, hsa-miR-183, hsa-miR-27a, hsa-miR-182, hsa-miR-25 | Positive | [47] |
| AGO3 | hsa-miR-30a, hsa-miR-21, hsa-miR-182, hsa-miR-30d, hsa-miR-378a, hsa-miR-30a, hsa-miR-183, hsa-miR-92a, hsa-miR-99b, hsa-miR-151a | Positive | [47] |

These studies provided a total of 104 experimentally validated functional miRNA-RBP combinations, which worked as a test to measure the effectiveness of the BNA in detecting them. Majority of these experimentally validated combinations were discovered by the BNA presented in the present study, bringing high confidence on the large number of novel miRNA and RBP functional combinations detected in the present study (>10,000 combinations were detected for mature miRNAs and RBP combinations).

A hierarchical cluster analysis was performed for clustering of miRNAs based on their expression data considering all the eight tissues together. miRNAs which expressed themselves in at least 50% samples were considered for clustering, resulting into a total of 632 miRNAs fulfilling the criteria. A total of 124 clusters were formed in which at least 80% RBPs were common for the associated corresponding miRNAs belonging to any given cluster. Out of 124 clusters, 96 clusters (75.9%) had more than 56% common pathways, 71% common biological process, and 73% common molecular functions between the miRNAs targets and RBPs. Fig 4 illustrates the cluster associations for 50 such miRNAs and their associated RBPs. This analysis further supported the functional miRNA and RBP associations captured through BNA.

**Fig 4.**
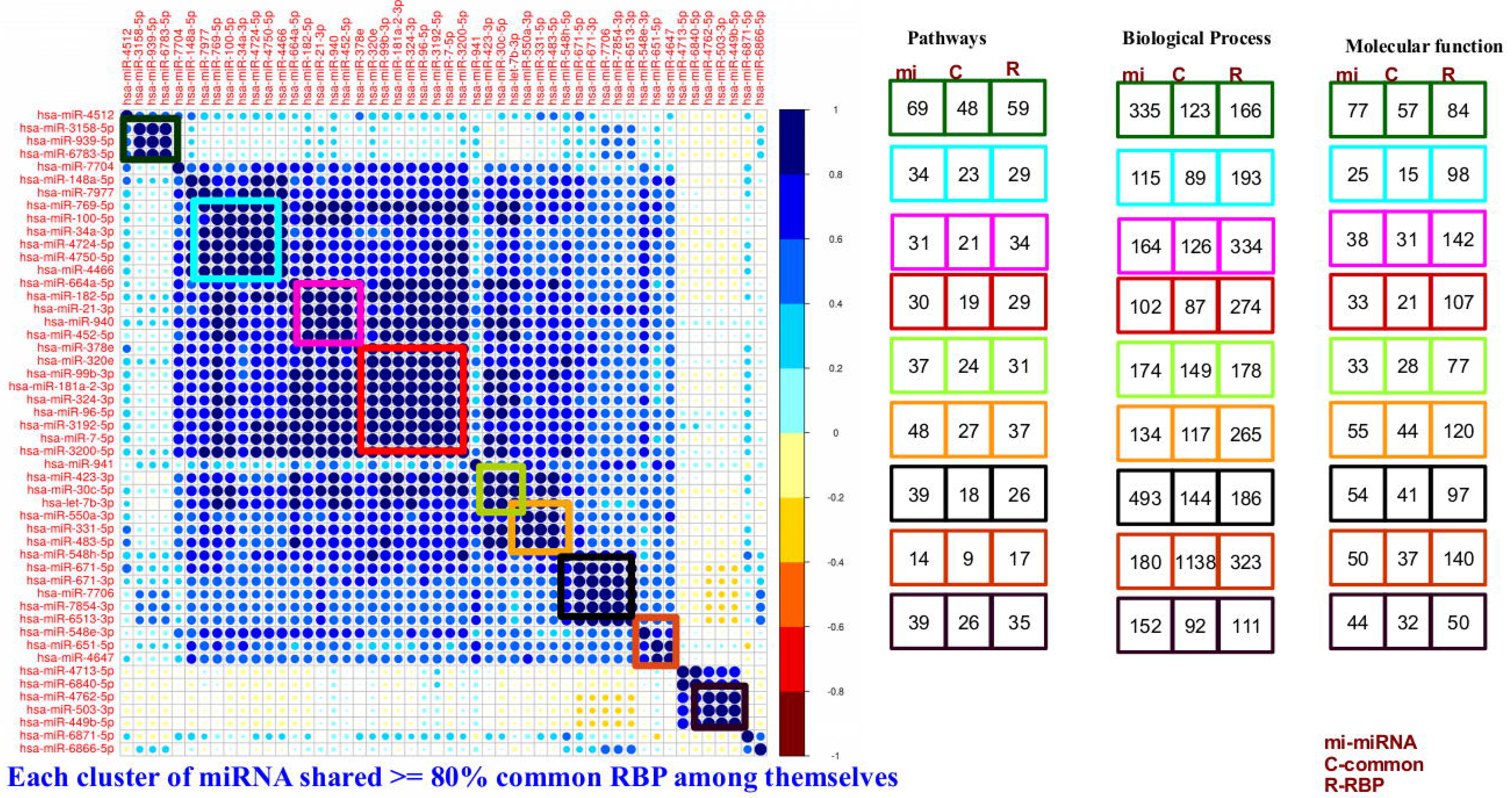
Correlogram based clustering identifies shared function by the cluster members. Commonly expressing miRNAs have common RBP partners, common control system, common biological roles. The miRNAs clustered in a common group were found to share more than 80% common RBPs which bound to their pre-miRNAs. This diagram represents different clusters of miRNAs along with the members in different cluster. The rectangle boxes display the common pathways, biological process and molecular functions between miRNAs and their associated RBPs belonging to same cluster.

A small test for functional unity between the miRNAs and their observed associated RBPs was done. The logic was that if the miRNAs were dependent upon these RBPs for their biogenesis and display functional similarity, then they would be able to distinguish the cell states similarly. Based on their normalized expression data, k-means clustering was performed to distinguish between lung cancer and normal condition cases with miRNAs expression data and associated RBPs expression data, separately (Fig 5). Similar results were obtained for both the clusterings where the two cell states were separated similarly. The experiment was repeated upon thyroid vs normal condition also and the same was observed here also: RBPs mirroring the result of associated miRNAs. This all underscored again that the BNA provided the functional associations between the RBPs and miRNAs.

**Fig 5.**
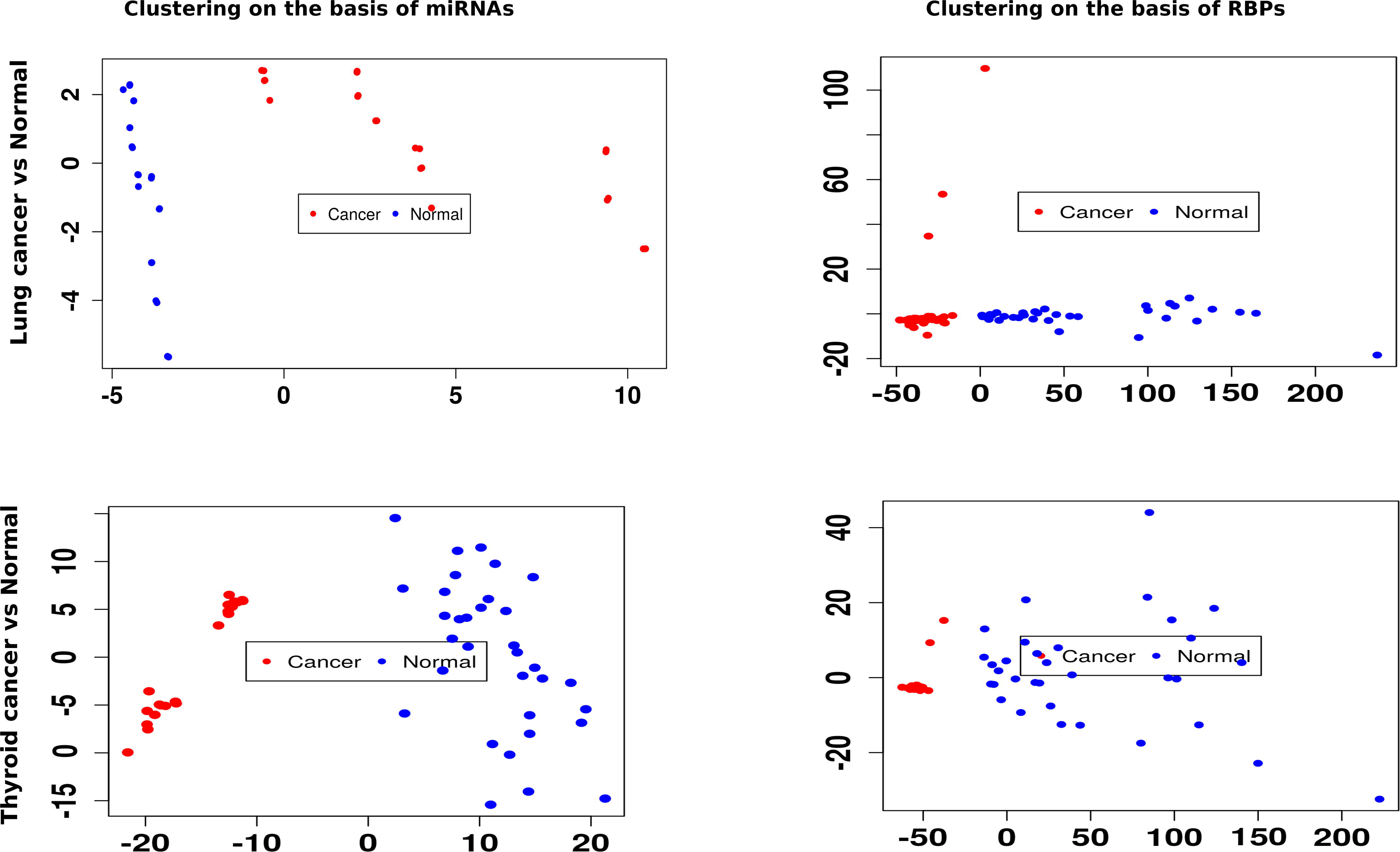
miRNAs and associated RBPs distinguish between cell lines similarly. This set of plots displays the discrimination between lungs cancer vs normal samples and thyroid cancer vs normal samples based on k-means clustering on miRNAs and associated RBPs, separately. The associated miRNAs and RBPs gave similar results, reflecting their functional association.

Besides revealing the most confident RBP and miRNA associations, the BNA also provided the causality nets for all such associations which may be considered as the logic units capable to decide the network flow towards miRNA biogenesis. In total, the RBP:miRNA networks reconstructed through BNA implicated ~1,400 unique genes as the members of these networks controlling the steps of miRNA biogenesis. Fig 6 illustrates the distributions of the top 30 most common genes across the various networks for miRNA biogenesis as well as total distribution patterns of the responsible networks’ genes. For any given miRNA’s network, there existed many paths to reason miRNA biogenesis in different conditions. However, few of these paths were dominant ones which were followed in several conditions and around which several conditions’ paths clustered with high degree of similarity to those dominant paths.

**Fig 6.**
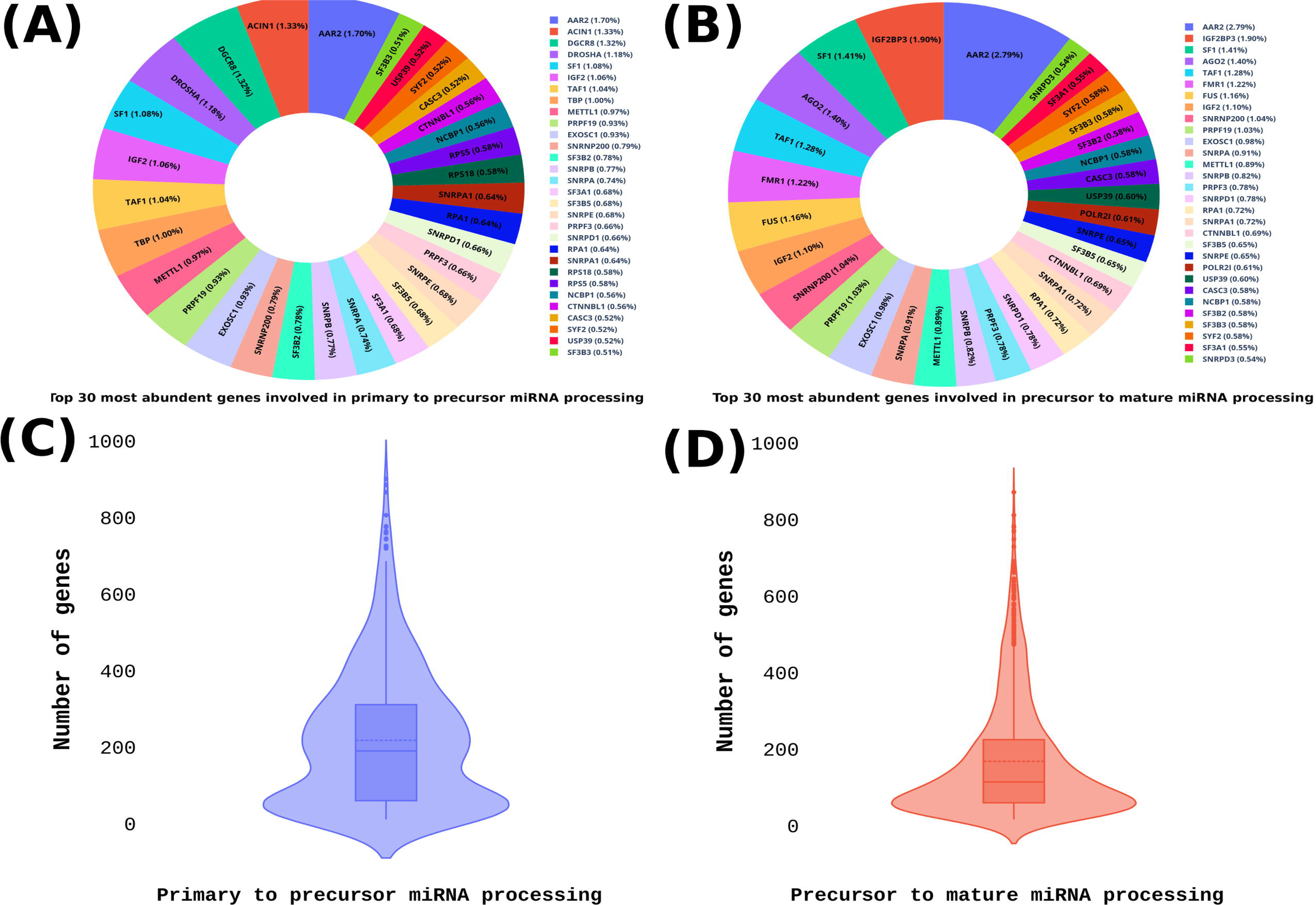
The most common genes across the conditional causal networks and distribution of genes across the networks. (A) The top 30 most common genes involved in the primary to precursor processing. (B) The top 30 most common genes involved in precursor to mature miRNA processing. The violin plots show the number of genes distribution across the identified networks involved in miRNA processing from primary to precursor (C), and precursor to mature miRNA(D).

### Machine learning on the Bayesian networks successfully predicts miRNA profiles

The BNA screened those RBPs from the CLIP-seq interaction data which displayed functional association with the bound miRNAs as well as it also extracted the directed networks which could define the causality for the associated miRNA biogenesis. It did one more interesting job which empowered the possibility to design a software system which could predict miRNA profiles. The network components discovered by the BNA may be seen as the feature variables holding the component genes set expression values which could work as the input vector to a machine learning system. Simultaneously, the associated miRNA’s expression data for any considered sample could be treated as the target value. This way, it was possible to train and build machine-learning models which could accurately predict the miRNA profiles even in the absence of miRNA profiling experiments like RNA-seq or arrays. Such tool becomes very important to work for conditions where miRNA profile is unknown or cost cutting on running miRNA profiling experiments is desired. More so, when large number of samples and conditions are to be studied.

To implement this, a machine learning approach, XGBoost regression, was used. The RNA-seq and miRNA-seq expression data were collected from TCGA database for seven different tissues such as bladder, brain, breast, cervix, esophagus, kidney and head-neck for model building and testing, as described in the methods section. Different types of machine learning regression techniques were tested and XGBoost regression emerged as the best one while scoring high consistency and an average accuracy of 91% (Accuracy range: 87% to 94%) for the modeled miRNAs (models for a total of 1,204 miRNAs), covering wide range of experimental conditions (total 30 experimental conditions) and tissue types (total 15 tissue types). The prediction rule was fine tuned by identifying the optimal combination of hyper-parameters that further minimized the objective function. With the optimal values of the hyper-parameters and number of trees, the regression was retrained and applied to the testing data to predict a new series of miRNA profiles and evaluate the accuracy of the model. The model accuracy was tested based on the RMSE (Root Mean Square Error) and RMAPE (Relative Mean Absolute Percentage Error). The 10-fold RMSE cross-validation was also found consistent when compared to the model RMSE which indicated the consistency of these models with highly reliable predictions. This also needs to be noted that in the present study, the amount of instances and samples were large enough to carry out robust measure like 10 folds cross-validation. However, if the sample size or the number of reported instances are small, one may opt for bootstrap re-sampling based approaches as bootstrapping is expected to perform good in such situation and give reliable result. Further to this, a t-test between the training and testing performance scores was found insignificant, reconfirming reliability of the trained models and their consistent performance on new and unseen data. Fig 7(A) presents a violin plot of these three performance metrics distribution for the testing and training datasets. The observed and predicted expression levels obtained from the XGBoost regression model for six miRNAs as examples is given in Fig 7(D).

**Fig 7.**
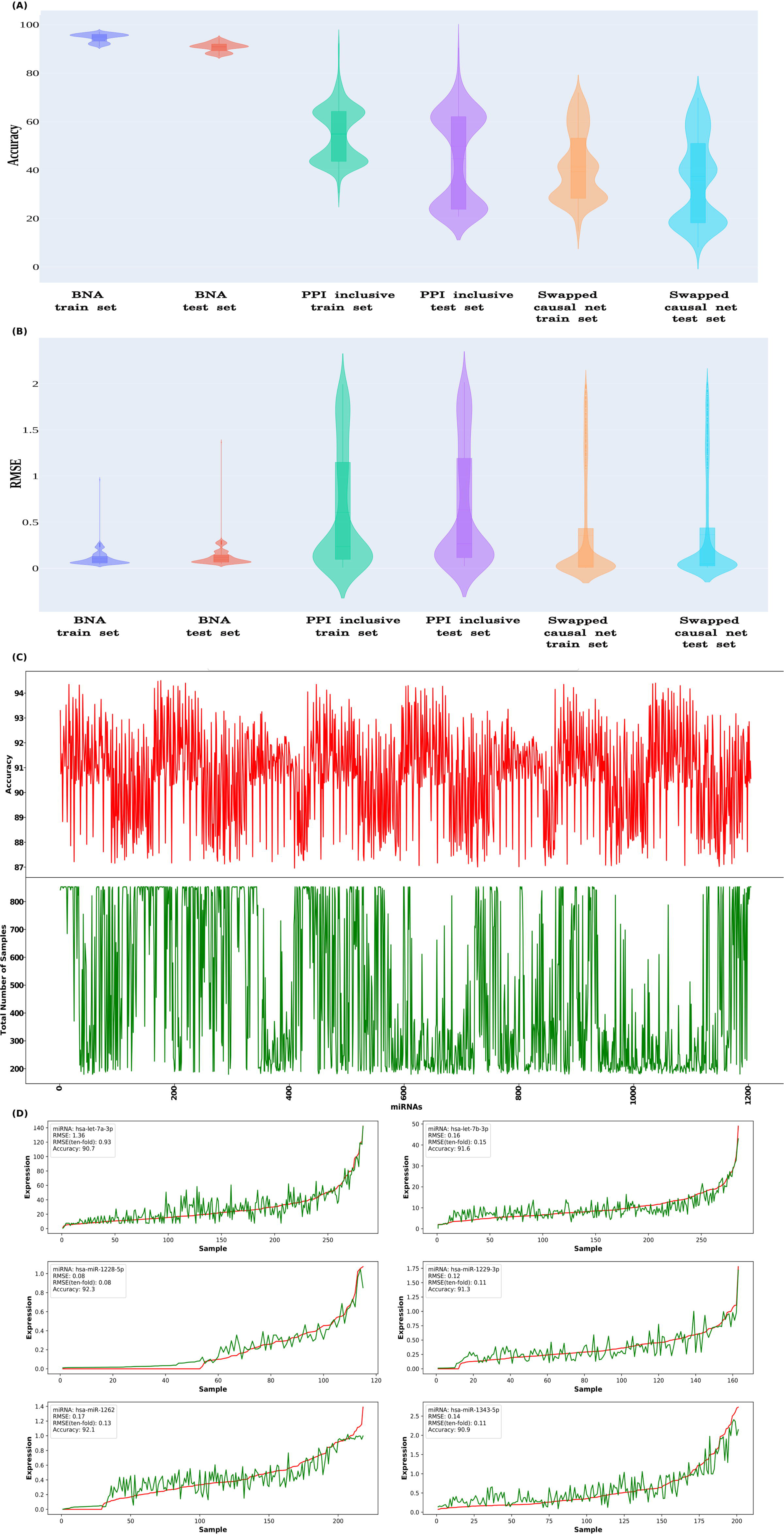
The causal network components’ expression data is enough to accurately predict miRNA profiles for any given condition. The miRNA expression levels were predicted from the RNA-seq expression data using XGBoost machine-learning. The models’ performance was measured based on accuracy and RMSE. The performance observed for the actual BNA causal nets based models were far superior than those models which used much bigger predecessor static networks (PPI-inclusive) from which causal nets were derived and models where the causal networks were randomly swapped to build the models with the causal net for some other miRNA (Swapped causal nets). Distribution of (A) Accuracy and (B) RMSE values for these models for all studied miRNAs is given in the form of violin plots. C) Accuracy observed for each miRNA when tested across the different experimental samples test set. The average accuracy was calculated over the 10 times sampled test sets, where every time the data was split in 70:30 ratio between training and test sets, and model was built from the scratch every time. The red colored plot shows the accuracy of the miRNAs for the given test sample set (the green colored counterpart). On x-axis 1,204 miRNAs are ordered in lexicographically order. Further details on samples and observed performance metrics are given in S6 Table, (D) The observed and predicted expression levels obtained for six representative miRNAs shows the high level of accuracy when studied across a large number of samples. The red line represents the actual expression observed for the miRNAs in the given samples, while the green line represents the predicted expression profile for the miRNAs.

Further to this, in order to evaluate that degree to which such high performance accuracy depended upon the identified causal network components and how specific they have been, a couple of tests was conducted. Since the BNA derived causal net’s genes for any given miRNA defined the feature sets for the machine learning model for the respective miRNAs, replacing these genes with some other genes into the machine learning model would help to measure the level of importance the BNA derived gene set held for the accurate assessment of the associated miRNA’s expression. Thus, in order to get a fare test done one would require to retrain and build a comparative model with the replaced feature set genes while mapping to the associated miRNA. The null hypothesis would suggest that no significant difference between the performance of the two compared models existed. In the first test the causal network genes defining the input feature for the associated miRNAs were replaced by the component genes which defined the initial network comprising all involved PPI partners also and providing much bigger super-set network than its corresponding Bayesian causal networks. For all the miRNAs, the corresponding XGBoost machine learning regression model was re-built using this much larger initial network input. The rebuilt models were tested for their capability to predict the corresponding miRNA’s expression profile as was done earlier for the identified causal net components for the associated miRNA. This test helped us to fathom that if a much bigger predecessor network lacking the causal information could perform better than the identified Bayesian causal nets for the associated miRNAs. Its comparison with Bayesian causal net based models clearly showcased that how the causal nets help in concentrating upon the relevant components to derive strong information and prediction system which is otherwise lost in the larger gene set with no causal information.

In the second test, the causal network component genes were swapped across the miRNAs and trainingof the machine learning system was done followed by performance testing in the similar manner as was done for the original models. This testing we called as the scrambled sets testing which tested if the actual causal network of any miRNA is replaced by the causal net identified for some other miRNAs gives the same level of performance as was observed for the original one with the actual associated causal net for the given miRNA.

For the first test the average accuracy for the training system could never breach the limit of 54.73%, and on the testing dataset it could touch an average accuracy of 44.67%, though the performance varied a lot across the miRNAs (Fig 7 (A,B)) (S6 Table). This clearly underscored that the derived causal networks found associated with miRNAs were specific and carried forward much information than their far bigger static network predecessor which did not capture causality. Similarly, in the second test the average accuracy dropped drastically to 41.83% for the training and 35.56% for the testing sets, respectively. Further, a t-test was performed between the BNA feature based model performance matrices (Accuracy, RMAPE and RMSE) and model performance metrics obtained during these two tests. The t-tests were found to be highly significant (p-value < 10^−48^) when compared to the BNA causal nets based models, suggesting significant difference in the observed performances of the models raised during these tests when compared with the actual BNA based models. In overall, these tests strongly underscored that the observed high accuracy of miRNA profile prediction with associated BNA causal net components was highly specific to the identified relationships between the causal nets and associated miRNAs. The detected causal relationships very aptly reflected themselves in making accurate miRNAs expression predictions which is not a random event.

Further level of testing was done for the developed models across 431 different samples covering different conditions which included different tissue types such as liver, prostrate, pancreas, ovary, lungs LAUD, lungs LUSC, adrenal gland ACC and adrenal gland PCPG, which were not covered in any of the studies taken in the section above. Out of 431 samples, 308 (72%) scored more than 90% accuracy, while 118 samples (27%) had more than 85% accuracy. S7 Table contains complete details of the performance measures on this data-set. The high degree of testing score consistency with high accuracy across such large number of experimental test data confirmed the developed tool as a highly reliable one and first of its kind to profile miRNAs in any given condition without any need to do miRNA sequencing or array profiling experiments. This need to be pointed out that in order to keep the entire study totally unbiased and without any chance of data-set driven memory, at every stage different data sources were taken and it was ensured that no overlap of any sort happened. This is why the data sources used to construct the Bayesian causal nets differed totally from the sources used to build the machine learning models, testing sets, and the further validation data-sets. All of them differed from each other in totality.

Fig 8 provides a final go through the basic work-flow of the prediction system implementation. From CLIP-seq data, most potential interacting miRNAs and RBP partners were extracted, which were screened further for relevant functional interactions while building upon the causality using BNA. For this, PPI data of all available protein-protein interactions were added to the identified interacting RBPs. The directionality and network flow were determined using available expression data for all these genes and associated miRNA from various experimental conditions. As can be seen from the figure, the flow of the network could follow many paths leading to miRNA biogenesis. Few of them were common to many conditions (thicker edges), while some work in antagonistic way (red lines). Such causal network works as the features vector for the XGBoost regression system where network gene components expression data worked as the input and associated miRNA expression data as the target to build the learning system first. The built model thereafter becomes capable to predict the miRNA profiles accurately for any condition from RNA-seq data itself without any need of running separate miRNA profiling experiments like miRNA-seq or array profiling etc.

**Fig 8.**
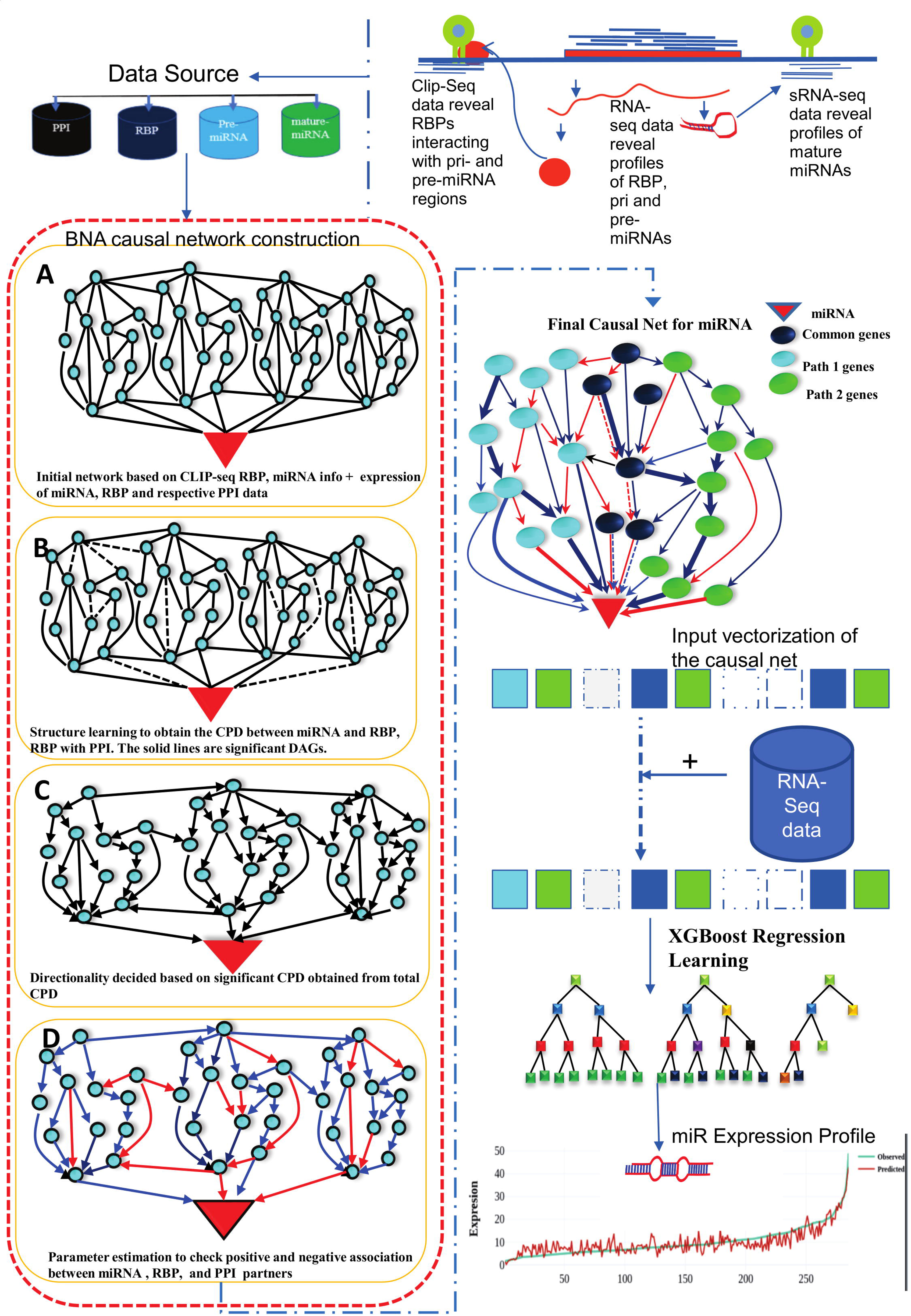
Work-flow of miRbiom prediction system implementation. High-throughput data from various platforms along with PPI data helped to build the initial network. BNA helped to reveal the functionally important connections and relationships, trimming the initial network while bringing directionality, causality, and preference. The fading edges represent insignificant associations, red edges are antagonist associations, and thickness of an edge is proportional to its recurrence/importance across various conditions. The final network for each miRNA works as an instruction set for the machine learning system (XGBoost) for learning and prediction system building. This uses RNA-seq data for the network components and miRNA expression data for various experimental conditions as the target to learn and build the prediction system. The finally built prediction system can accurately predict miRNA profiles for wide range of conditions.

The miRNA profile prediction system has been implemented as a webserver at https://scbb.ihbt.res.in/miRbiom-webserver/. The source code along with the model files for 1,204 miRNAs to run the standalone miRNAs profiling and benchmarking parts have been made available at https://scbb.ihbt.res.in/miRbiom-webserver/SC/Source_code_miRbiom-v-1.0.tar.gz. The user needs to provide the RNA-seq or any expression profiling experiment data for any given condition. This data is run through the trained XGBoosting models which generate a relative expression scores for various miRNAs capturing the potential expression profile of the miRNAs for the given condition. It generates a plot of expression profiles of various miRNAs in an interactive form. For validation purpose, a benchmarking tab is also provided in the result page where the user can provide their actual experimental profiling data for the miRNAs and compare the predicted profile. Selections can be made here to study the miRNA targets for their functional enrichment as well as pathways analysis in interactive and visually rich manner. miRNA target information from various databases like miRTarbase, TarBase, and Targetscan has been provided. Provisions have also been made to map the miRNA targets in a collective manner and view them in KEGG pathways maps. The server has been implemented using D3JS visualization library, Python, Javascript, PHP, and HTML5. Enrichment analysis was implemented through Enrichr [34] in the back-end. Fig 9 provides an overview of the miRbiom webserver.

**Fig 9.**
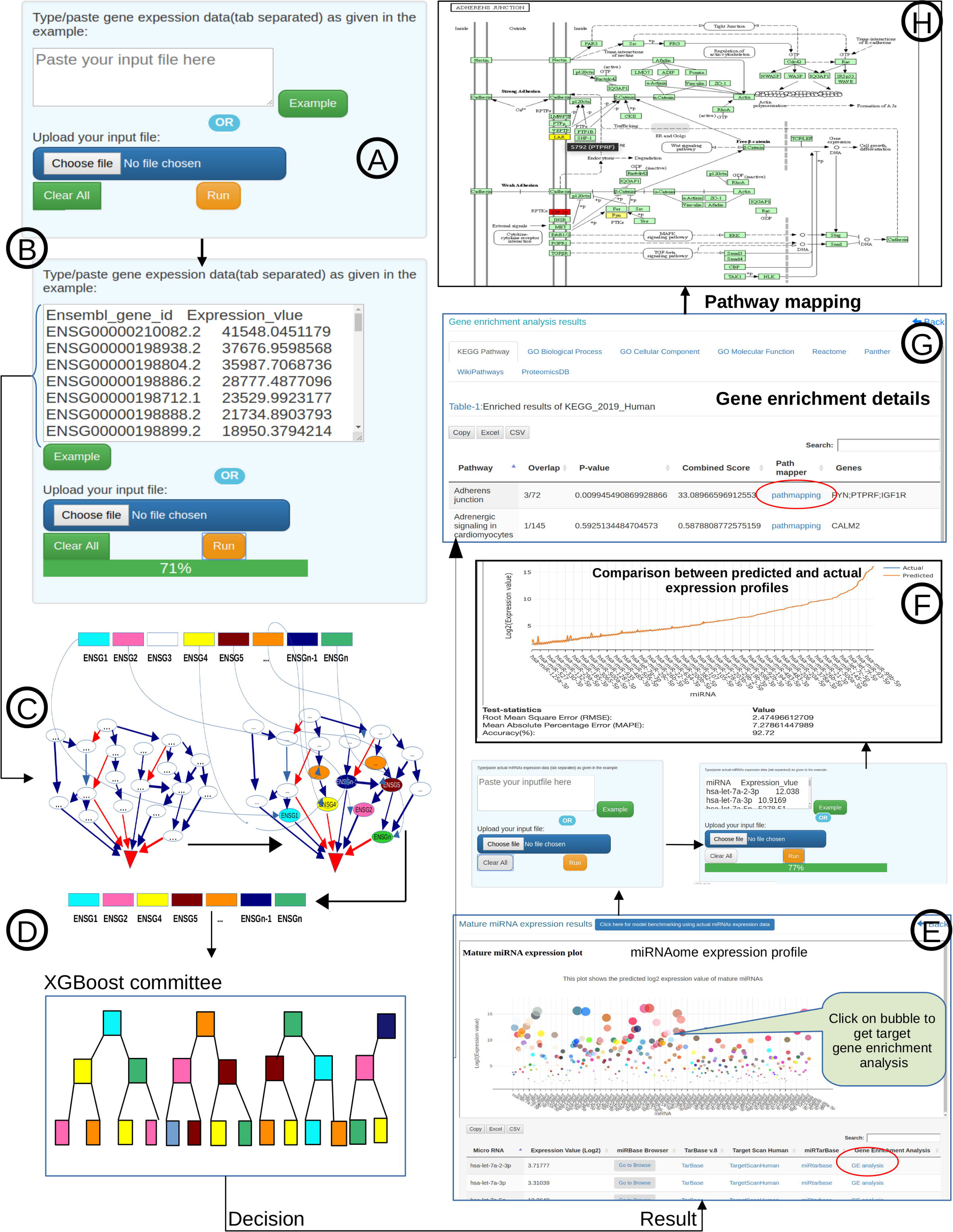
miRbiom webserver implementation. A) Input data box where the user can either paste or load the input file, B) Input is transcriptome expression data with Ensembl IDs which is in turn normalized by the server, C) The causal network components are consulted and loaded with their respective expression data, D) XGBoost regression uses the network expression data to generate a jury of classifiers which commonly reach to decision with the expression profiles for various miRNAs in one go, E) Expression profile result is generated in the form interactive plot and table, (F) A module is provided to compare the predicted profile with the actual profile for test purpose, if the user wishes to, G) Functional enrichment analysis tabs for the targets of the miRNAs, H) Pathways mapping module for the miRNA targets.

## Conclusion

In the present study, the causal nets for miRNA biogenesis were identified while leveraging from a wide range of high-throughput data sources, experimental conditions, and cell lines. The causal nets were able to report much larger set of functional miRNAs and RBP interactions which were validated through some of already existing experimental data. The final step of this study not just re-validated the approach, but clearly displayed how much successful it could be in correctly predicting the miRNA expression profile while depending upon the found causal nets when machine-learning training is done on them. The networks worked as the training modules to raise a machine-learning system where the user just needs to provide the RNA-seq or any transcriptome profiling data which is mapped to the identified causal nets. Various paths of these causal nets are explored to finally take the decision on the profile of the miRNAs for the given experimental condition. In doing this all, this entire software, miRbiom, also democratizes the process of miRNA profiling itself, as it provides freedom to directly and accurately predict the miRNA expression profile from the transcriptome data of experiments like RNA-seq/arrays without any need to carry out miRNA expression profiling experiments like miRNA-seq etc separately.

From here now, we expect that more emphasis is given on the studies focusing on the relationship between the two post-transcriptional regulators, miRNAs and RBPs, and their associations. Availability of more high-throughput data from CLIP-seq, RNA-seq, and miRNA-seq will be immensely helpful to develop models for more miRNAs and achieve larger universality. The developed software may be extended further to other species given the above mentioned high-throughput data for them are also generated. Further to this, consideration of RNA modifications and epitranscriptomics data may mark the new level and bring us more closer to the system level maps for miRNA biogenesis.

## Declarations

### Availability of data and materials

All the secondary data used in this study were publicly available and their due references and sources have been provided. All data and information generated/used, methodology related details etc have also been made available in the supplementary data files provided along with and also made available through the related open access server at https://scbb.ihbt.res.in/miRbiom-webserver/. The standalone version of the program has also been made available at Github https://github.com/SCBB-LAB/miRbiom.

### Authors’ contributions

UKP and NKS carried out the computational and statistical part of the study. PK developed the web-server. AK helped in computational analysis. SG contributed in performance measure study. RS conceptualized, designed, analyzed and supervised the entire study. UKP, NKS, and RS wrote the MS.

## Acknowledgments

We are thankful to the Director, CSIR-IHBT for his kind support in getting the experimental validation work carried out at CSIR-IHBT. We are thankful to Dr. Dharam Singh for his discussions. UKP and PK are thankful to ICAR, New Delhi for providing support in Ph.D. UKP, NKS, and PK are thankful to Academy of Scientific and Innovative Research (AcSIR) for their Ph.D. enrollment. We are also thankful to Mr. Anoop Kumar assisting in the MS preparation. This MS has CSIR-IHBT MSID # 4643.

## Ethics approval and consent to participate

Not applicable.

## Supporting information

**S1 Table. Detailed description of CLIP-seq datasets used in this study for different RBPs.** This table contained the accession id, study id, different clip-seq variants and different experimental conditions of each RBP.

**S2 Table. Detailed description of RNA-seq and sRNA-seq expression data collected from TCGA database.**

**S3 Table. Detail of 431 samples collected from TCGA used for validation**

**S4 Table. This table described number of Pri-miRNA and Pre-miRNA having binding site on each RBP out of 1,881 miRNAs.**

**S5 Table. This table describes number of RBPs have binding sites in each Pri-miRNA and Pre-miRNA.**

**S6 Table. Performance comparison among BNA based features, all initial input data for BNA(PPI) and shuffled feature set in terms of Accuracy, RMAPE and RMSE.**

**S7 Table. Accuracy, RMAPE and RMSE values for validation set of 431 samples**

## References

1. Lewis BP, Shih I-hung, Jones-Rhoades MW, Bartel DP, Burge CB. Prediction of mammalian microRNA targets. Cell. 2003;115: 787–798. doi:10.1016/s0092-8674(03)01018-3

2. Kozomara A, Birgaoanu M, Griffiths-Jones S. miRBase: from microRNA sequences to function. Nucleic Acids Research. 2019;47: D155–D162. doi:10.1093/nar/gky1141

3. Michlewski G, Cáceres JF. Post-transcriptional control of miRNA biogenesis. RNA. 2019;25: 1–16. doi:10.1261/rna.068692.118

4. Siomi H, Siomi MC. Posttranscriptional regulation of microRNA biogenesis in animals. Mol Cell. 2010;38: 323–332. doi:10.1016/j.molcel.2010.03.013

5. Ratnadiwakara M, Mohenska M, Änkö M-L. Splicing factors as regulators of miRNA biogenesis - links to human disease. Semin Cell Dev Biol. 2018;79: 113–122. doi:10.1016/j.semcdb.2017.10.008

6. Kim Y-K, Kim B, Kim VN. Re-evaluation of the roles of DROSHA, Export in 5, and DICER in microRNA biogenesis. Proc Natl Acad Sci U S A. 2016;113: E1881–1889. doi:10.1073/pnas.1602532113

7. Newman MA, Thomson JM, Hammond SM. Lin-28 interaction with the Let-7 precursor loop mediates regulated microRNA processing. RNA. 2008;14: 1539–1549. doi:10.1261/rna.1155108

8. Morlando M, Dini Modigliani S, Torrelli G, Rosa A, Di Carlo V, Caffarelli E, et al. FUS stimulates microRNA biogenesis by facilitating co-transcriptional Drosha recruitment. EMBO J. 2012;31: 4502–4510. doi:10.1038/emboj.2012.319

9. Zisoulis DG, Kai ZS, Chang RK, Pasquinelli AE. Autoregulation of microRNA biogenesis by let-7 and Argonaute. Nature. 2012;486: 541–544. doi:10.1038/nature11134

10. Westholm JO, Lai EC. Mirtrons: microRNA biogenesis via splicing. Biochimie. 2011;93: 1897–1904. doi:10.1016/j.biochi.2011.06.017

11. Jiang P, Coller H. Functional interactions between microRNAs and RNA binding proteins. Microrna. 2012;1: 70–79. doi:10.2174/2211536611201010070

12. Jha A, Mehra M, Shankar R. The regulatory epicenter of miRNAs. J Biosci. 2011;36: 621– 638. doi:10.1007/s12038-011-9109-y

13. Corley M, Burns MC, Yeo GW. How RNA-Binding Proteins Interact with RNA: Molecules and Mechanisms. Mol Cell. 2020;78: 9–29. doi:10.1016/j.molcel.2020.03.011

14. Treiber T, Treiber N, Plessmann U, Harlander S, Daiß J-L, Eichner N, et al. A Compendium of RNA-Binding Proteins that Regulate MicroRNA Biogenesis. Mol Cell. 2017;66: 270–284.e13. doi:10.1016/j.molcel.2017.03.014

15. Nussbacher JK, Yeo GW. Systematic Discovery of RNA Binding Proteins that Regulate MicroRNA Levels. Mol Cell. 2018;69: 1005–1016.e7. doi:10.1016/j.molcel.2018.02.012

16. Gahlan P, Singh HR, Shankar R, Sharma N, Kumari A, Chawla V, et al. De novo sequencing and characterization of Picrorhiza kurrooa transcriptome at two temperatures showed major transcriptome adjustments. BMC Genomics. 2012;13: 126. doi:10.1186/1471-2164-13-126

17. Bolger AM, Lohse M, Usadel B. Trimmomatic: a flexible trimmer for Illumina sequence data. Bioinformatics. 2014;30: 2114–2120. doi:10.1093/bioinformatics/btu170

18. Jiang H, Wong WH. SeqMap: mapping massive amount of oligonucleotides to the genome. Bioinformatics. 2008;24: 2395–2396. doi:10.1093/bioinformatics/btn429

19. Salzman J, Jiang H, Wong WH. Statistical Modeling of RNA-Seq Data. Stat Sci. 2011;26. doi:10.1214/10-STS343

20. Friedländer MR, Mackowiak SD, Li N, Chen W, Rajewsky N. miRDeep2 accurately identifies known and hundreds of novel microRNA genes in seven animal clades. Nucleic Acids Res. 2012;40: 37–52. doi:10.1093/nar/gkr688

21. Ritchie ME, Phipson B, Wu D, Hu Y, Law CW, Shi W, et al. limma powers differential expression analyses for RNA-sequencing and microarray studies. Nucleic Acids Res. 2015;43: e47. doi:10.1093/nar/gkv007

22. Uren PJ, Bahrami-Samani E, Burns SC, Qiao M, Karginov FV, Hodges E, et al. Site identification in high-throughput RNA-protein interaction data. Bioinformatics. 2012;28: 3013–3020. doi:10.1093/bioinformatics/bts569

23. Saini HK, Enright AJ, Griffiths-Jones S. Annotation of mammalian primary microRNAs. BMC Genomics. 2008;9: 564. doi:10.1186/1471-2164-9-564

24. Szklarczyk D, Franceschini A, Wyder S, Forslund K, Heller D, Huerta-Cepas J, et al. STRING v10: protein-protein interaction networks, integrated over the tree of life. Nucleic Acids Res. 2015;43: D447–452. doi:10.1093/nar/gku1003

25. Anderson JC, Gerbing DW. Structural Equation Modeling in Practice: A Review and Recommended Two-Step Approach. PSYCHOLBULL. 1988;103: 411–423. doi:10.1037/0033-2909.103.3.411

26. Gupta S, Kim HW. Linking structural equation modeling to Bayesian networks: Decision support for customer retention in virtual communities. European Journal of Operational Research. 2008;190: 818–833. doi:10.1016/j.ejor.2007.05.054

27. Beck A, Tetruashvili L. On the Convergence of Block Coordinate Descent Type Methods. SIAM J Optim. 2013;23: 2037–2060. doi:10.1137/120887679

28. Aragam B, Zhou Q. Concave Penalized Estimation of Sparse Gaussian Bayesian Networks. Journal of Machine Learning Research. 2015;16: 2273–2328.

29. Zhang C-H. Nearly unbiased variable selection under minimax concave penalty. The Annals of Statistics. 2010;38: 894–942. doi:10.1214/09-AOS729

30. Schober P, Boer C, Schwarte LA. Correlation Coefficients: Appropriate Use and Interpretation. Anesth Analg. 2018;126: 1763–1768. doi:10.1213/ANE.0000000000002864

31. Panwar B, Omenn GS, Guan Y. miRmine: a database of human miRNA expression profiles. Bioinformatics. 2017;33: 1554–1560. doi:10.1093/bioinformatics/btx019

32. Lachmann A, Torre D, Keenan AB, Jagodnik KM, Lee HJ, Wang L, et al. Massive mining of publicly available RNA-seq data from human and mouse. Nat Commun. 2018;9: 1366. doi:10.1038/s41467-018-03751-6

33. Parkinson H, Kapushesky M, Shojatalab M, Abeygunawardena N, Coulson R, Farne A, et al. ArrayExpress--a public database of microarray experiments and gene expression profiles. Nucleic Acids Res. 2007;35: D747–750. doi:10.1093/nar/gkl995

34. Kuleshov MV, Jones MR, Rouillard AD, Fernandez NF, Duan Q, Wang Z, et al. Enrichr: a comprehensive gene set enrichment analysis web server 2016 update. Nucleic Acids Res. 2016;44: W90–97. doi:10.1093/nar/gkw377

35. Huang H-Y, Lin Y-C-D, Li J, Huang K-Y, Shrestha S, Hong H-C, et al. miRTarBase 2020: updates to the experimentally validated microRNA-target interaction database. Nucleic Acids Res. 2020;48: D148–D154. doi:10.1093/nar/gkz896

36. Chen T, Guestrin C. XGBoost: A Scalable Tree Boosting System. Proceedings of the 22nd ACM SIGKDD International Conference on Knowledge Discovery and Data Mining. 2016; 785–794. doi:10.1145/2939672.2939785

37. Ramalingam P, Palanichamy JK, Singh A, Das P, Bhagat M, Kassab MA, et al. Biogenesis of intronic miRNAs located in clusters by independent transcription and alternative splicing. RNA. 2014;20: 76–87. doi:10.1261/rna.041814.113

38. Powers S, DeJongh M, Best AA, Tintle NL. Cautions about the reliability of pairwise gene correlations based on expression data. Front Microbiol. 2015;6: 650. doi:10.3389/fmicb.2015.00650

39. Kooshapur H, Choudhury NR, Simon B, Mühlbauer M, Jussupow A, Fernandez N, et al. Structural basis for terminal loop recognition and stimulation of pri-miRNA-18a processing by hnRNP A1. Nat Commun. 2018;9: 2479. doi:10.1038/s41467-018-04871-9

40. Alarcón CR, Goodarzi H, Lee H, Liu X, Tavazoie S, Tavazoie SF. HNRNPA2B1 Is a Mediator of m(6)A-Dependent Nuclear RNA Processing Events. Cell. 2015;162: 1299–1308. doi:10.1016/j.cell.2015.08.011

41. Wu H, Sun S, Tu K, Gao Y, Xie B, Krainer AR, et al. A splicing-independent function of SF2/ASF in microRNA processing. Mol Cell. 2010;38: 67–77. doi:10.1016/j.molcel.2010.02.021

42. Zhao L, Mao Y, Zhao Y, He Y. DDX3X promotes the biogenesis of a subset of miRNAs and the potential roles they played in cancer development. Sci Rep. 2016;6: 32739. doi:10.1038/srep32739

43. Abdelmohsen K, Tominaga-Yamanaka K, Srikantan S, Yoon J-H, Kang M-J, Gorospe M. RNA-binding protein AUF1 represses Dicer expression. Nucleic Acids Res. 2012;40: 11531–11544. doi:10.1093/nar/gks930

44. Hubé F, Ulveling D, Sureau A, Forveille S, Francastel C. Short intron-derived ncRNAs. Nucleic Acids Res. 2017;45: 4768–4781. doi:10.1093/nar/gkw1341

45. Li H-K, Mai R-T, Huang H-D, Chou C-H, Chang Y-A, Chang Y-W, et al. DDX3 Represses Stemness by Epigenetically Modulating Tumor-suppressive miRNAs in Hepatocellular Carcinoma. Sci Rep. 2016;6: 28637. doi:10.1038/srep28637

46. Scott MS, Avolio F, Ono M, Lamond AI, Barton GJ. Human miRNA precursors with box H/ACA snoRNA features. PLoS Comput Biol. 2009;5: e1000507. doi:10.1371/journal.pcbi.1000507

47. Dueck A, Ziegler C, Eichner A, Berezikov E, Meister G. microRNAs associated with the different human Argonaute proteins. Nucleic Acids Res. 2012;40: 9850–9862. doi:10.1093/nar/gks705

